# SpatialPPI 2.0: Enhancing Protein-Protein Interaction Prediction through Inter-Residue Analysis in Graph Attention Networks

**DOI:** 10.1101/2024.10.25.620355

**Authors:** Wenxing Hu, Masahito Ohue

## Abstract

Protein-protein interactions (PPIs) are fundamental to cellular functions, and accurate prediction of these interactions is crucial to understanding biological mechanisms and facilitating drug discovery. SpatialPPI 2.0 is an advanced graph neural network-based model that predicts PPIs by utilizing interresidue contact maps derived from both structural and sequence data. By leveraging the comprehensive PINDER dataset, which includes interaction data from the RCSB PDB and the AlphaFold database, SpatialPPI 2.0 improves the specificity and robustness of the prediction of PPI. Unlike the original SpatialPPI, the updated version employs interaction interface prediction as an intermediate step, allowing for a more effective assessment of interactions between isolated proteins. The model utilizes Graph Attention Networks (GAT) and Graph Convolutional Networks (GCN) to capture both local and global structural features. SpatialPPI 2.0 outperforms several state-of-the-art PPI and interface predictors, demonstrating superior accuracy and reliability. Furthermore, the model shows robustness when using structures predicted by AlphaFold, indicating its potential to predict interactions for proteins without experimentally determined structures. SpatialPPI 2.0 offers a promising solution for the accurate prediction of PPIs, providing insight into protein function and supporting advances in drug discovery and synthetic biology. SpatialPPI 2.0 is available at https://github.com/ohuelab/SpatialPPI2.0

## I. Introduction

Predicting protein-protein interactions (PPIs) is a fundamental problem in structural biology, as these interactions are at the core of many biological processes and pathways. PPIs play a vital role in cellular functions such as signal transduction, immune response, and metabolic regulation, which are crucial to maintaining cellular homeostasis. Understanding PPIs enables insight into cellular machinery and biological networks, thereby helping in the identification of novel drug targets and the development of therapeutic strategies. PPI prediction is of particular importance for understanding disease mechanisms and designing intervention approaches. Moreover, accurate PPI predictions have significant applications in synthetic biology and protein engineering, where targeted manipulations of biological systems are required [1]–[3].

PPI prediction can be broadly categorized into sequencebased and structure-based approaches. Sequence-based methods are widely used due to the abundance of protein sequence data and their computational efficiency [4]–[6]. Sequencebased approaches, such as D-script [7] and Topsy-Turvy [8], exploit amino acid sequences and evolutionary information to predict interactions. D-script leverages neural networks to represent and predict PPIs by learning sequence patterns, while Topsy-Turvy uses transfer learning techniques to enhance the prediction performance based on sequence data. However, sequence-based approaches lack the spatial information of proteins, which is crucial in determining how proteins physically interact [9]. This limitation often restricts their ability to accurately distinguish between interacting and non-interacting proteins.

Structure-based PPI prediction methods, on the other hand, incorporate three-dimensional (3D) structural information of proteins, providing a more direct representation of their potential interaction [10], [11]. Structure-based methods can better predict the physical compatibility of binding surfaces and identify specific contact regions, which is an advantage over sequence-based approaches [12], [13]. Struct2Graph [14] and GNN-PPI [15] are two notable structure-based prediction methods. Struct2Graph converts 3D protein structures into graph representations and applies graph-based learning to infer interactions. GNN-PPI also employs graph neural networks to capture both local and global structural features of proteins. Although structure-based predictions offer greater specificity in identifying binding interfaces, their applicability is often hindered by the scarcity of experimentally determined protein structures and the computational expense associated with structural data processing [16]. Despite these challenges, structure-based methods provide superior interpretability, making them more desirable when structural data are available. Consequently, we chose to adopt a structure-based prediction strategy for this study.

The prediction of interaction interfaces is a critical aspect of structure-based PPI prediction [17]–[20]. Interaction interfaces are the specific regions of the protein surface that are involved in binding interactions. Methods such as DeepInter [21] and CDPred [22] focus on predicting these interaction interfaces. DeepInter uses graph neural networks to predict inter-residue contacts, while CDPred employs homology-based approaches to identify probable interface residues. Both DeepInter and CDPred rely heavily on UniProt Reference Clusters (UniRef) and homology searches [24], which may lead to limitations when dealing with novel proteins without homologous sequences [25]. However, predicting interaction interfaces as an intermediate step offers significant benefits in overall PPI prediction, as it allows models to focus on specific regions of the protein structure rather than considering the entire molecule.

A key aspect of our study is the utilization of the PINDER dataset (Protein Interaction Dataset and Evaluation Resource) [26], which provides a comprehensive collection of protein-protein interaction and docking data. Compared to other state-of-the-art datasets such as DIPS-Plus [27], ProteinFlow [28], and PPIRef [29], PINDER contains more than 500 times more data and offers greater diversity to train flexible models capable of adapting to predicted structures. PINDER includes structural data from both the RCSB Protein Data Bank (PDB) [30] and the AlphaFold Database [31], employing Interface-Based Clustering and Deleaking techniques to ensure high-quality, non-redundant data. The large size and quality of the PINDER dataset make it an excellent resource for training models to predict both protein interfaces and overall interactions, thus enhancing model robustness and generalizability. The training set of PINDER comprises millions of dimers, offering diverse examples that can facilitate the model’s capability to accurately capture different types of interactions, including challenging cases involving diverse protein structures.

Recent advances in protein structure prediction, such as AlphaFold3 [32] and the AlphaFold Database [33], have provided valuable solutions to address the scarcity of protein structural data. AlphaFold represents a breakthrough in accurately predicting protein structures, even for those lacking experimental characterization [34]. However, AlphaFold’s predictions are not always reliable in distinguishing whether the input proteins can physically interact. AlphaFold has been observed to incorrectly predict well-folded complexes for protein pairs that do not interact [35]. This limitation complicates the direct application of AlphaFold to PPI prediction. However, analyzing the predicted structures by AlphaFold can provide valuable insight into potential PPIs.

To address these challenges, we present SpatialPPI 2.0, a novel structure-based PPI prediction framework that uses predicted protein-protein interaction interfaces as an intermediate result for more accurate PPI prediction. Compared to the original SpatialPPI [36], SpatialPPI 2.0 introduces a crucial enhancement by predicting interaction interfaces as intermediate steps, significantly improving its ability to assess interactions between isolated proteins. SpatialPPI 2.0 leverages graph attention networks (GATs) [37] and graph convolutional networks (GCNs) [38] to capture both local contact information and global structural features, providing an improved representation of protein complexes. By utilizing interaction interface predictions, our model is capable of enhancing the specificity of PPI predictions while maintaining robustness, even when relying on predicted structures. This study aims to demonstrate the utility of SpatialPPI 2.0 in effectively predicting PPIs and to highlight its applications in studying interactions involving proteins for which no experimental structures are available, thereby paving the way for advancements in drug discovery and synthetic biology.

## II. Methods and Materials

SpatialPPI 2.0 is a graph neural network-based deep learning model designed to predict protein-protein interactions by estimating inter-residue contact maps between two proteins, using both their structural and sequence information. The workflow is depicted in Fig. 1. Protein structure files (PDB) are processed to extract features, including amino acid sequence features and adjacency matrices, to construct graph representations of the proteins. The Interface Predictor then analyzes these graph features to predict the contact map between the two proteins. Based on this predicted contact map, the graph representation of the proteins is updated, and the Interaction Predictor is used to estimate the likelihood of interaction between the two proteins.

**Fig. 1.**
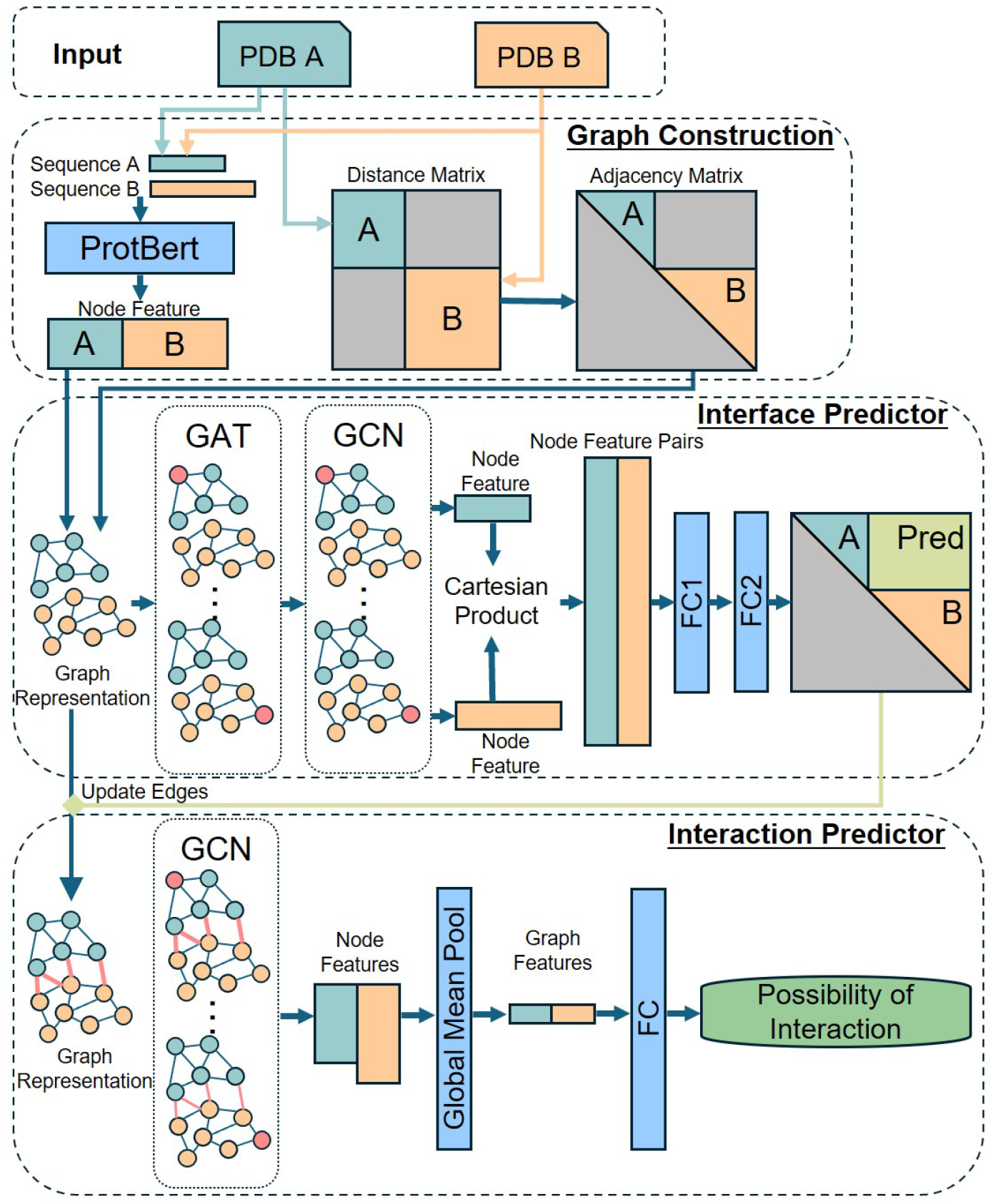
Detailed workflow of SpatialPPI 2.0. Input protein structures (PDB files) are used to extract both structural and sequence features. These features are represented as graph-based data, which are then processed using a combination of graph attention networks (GATs) and graph convolutional networks (GCNs) to predict inter-residue contacts. The predicted contact map is subsequently used to refine the graph representations, which are ultimately fed into an interaction predictor to determine the likelihood of a protein-protein interaction.

### A. Dataset Construction

The construction of a high-quality dataset is crucial for the performance and generalizability of deep neural network models. PINDER (Protein Interaction Dataset and Evaluation Resource) provides a comprehensive collection of proteinprotein interaction and docking data. PINDER includes structural data from both the RCSB PDB and AlphaFold Database and employs Interface-Based Clustering and Deleaking. Two interfaces are marked as similar if they meet the following criteria: iRMS *<* 5.0, IS-score *>* 0.3, log(P-value) *<* − 9.0, where iRMS is the interface RMSD after alignment, IS-score represents the Interface Similarity score, and log(P-value) is the logarithm of the statistical significance of the IS-score, determined from the distribution of interface scores using the iAlign methodology [39]. Based on the clustering results, the dataset is divided into training, validation, and test sets, with redundancy removed and filtering applied to ensure the highest quality. The training set of PINDER consists of 1,560,682 dimers from 42,220 clusters, of which 136,498 have paired apo structures and 566,171 have paired AlphaFold Database structures. We further filtered the dataset for our purposes. Pairs of residues with a C_*α*_ distance less than 8 Å are considered to be in contact [40]. Only residue pairs with more than eight contacts were used, and protein sequence lengths were restricted to between 35 and 300 residues. The data from PINDER were used to train both the protein interface prediction model and as a positive dataset for protein interaction prediction.

For non-interacting protein pairs, Negatome 2.0 [41] served as an important resource. Negatome 2.0 provides manually curated, experimentally verified non-interacting protein pairs, which are reliable for evaluating PPI prediction performance [42]. However, because of the limited availability of experimentally verified negative samples, we selected proteins from the PINDER test set, which had determined structures, as the negative test dataset for PPI prediction. The negative training dataset was generated by random sampling to create an equal number of negative samples as positive and was cross-checked with BioGRID [43] for validation. Table I summarizes all the datasets used in this study and their final filtered sizes.

**TABLE I.**
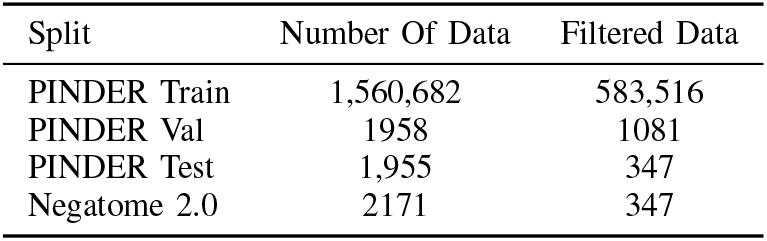
Datasets used in this study.

### B. Feature Extraction

The input data for the network are transformed into graph representations for processing by the neural network. Each residue in the protein is represented as a node in the graph, with features computed using the ProtBert model from ProtTrans [44]. Each residue is transformed into a 1024-dimensional vector representation. Compared to one-hot encoding, protein representations based on language models provide richer contextual information [45]. Moreover, using language models significantly improves the processing speed compared to database-based methods, such as retrieving embeddings from UniRef [46]. The language model also exhibits superior performance in predicting novel or designed proteins lacking homologous structures [47].

The C_*α*_ distance between the residues in the protein structure is used to compute a distance matrix, as shown in Fig. 2(a). The upper right section of the distance matrix, computed from the combined structure of two proteins, serves as the ground truth for the interface between the two proteins. Distance matrices computed separately for each protein are used as adjacency matrices for graph construction, as shown in Fig. 2(b). An edge is constructed between two residue nodes if the C_*α*_ distance between them is less than 8 Å, with the edge weight set to the distance between the residues. Based on the extracted residue nodes and adjacency matrix, the graph representation of the protein is generated and used as input to the neural network.

**Fig. 2.**
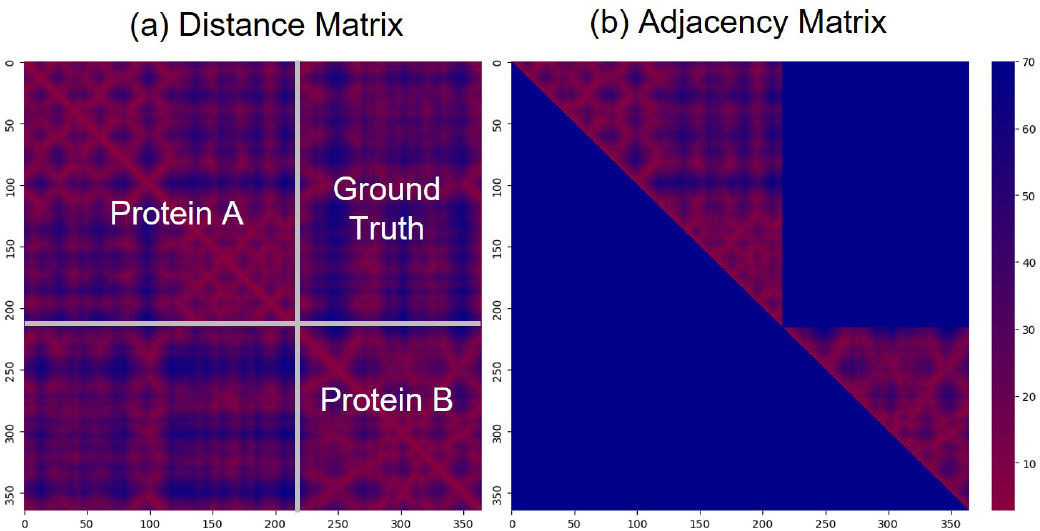
Feature extraction process for SpatialPPI 2.0. (a) The distance matrix is computed from the combined structure of Protein A and Protein B, with the upper right section representing the ground truth for the interface. (b) The adjacency matrix derived from the distance matrices of each protein is used for graph construction where edges represent residue-residue proximity.

### C. Interface Predictor

The Interface Predictor consists of a GAT layer and a GCN layer to calculate pairwise inter-residue features. GAT dynamically adjusts the weights of information propagation between nodes based on their features, allowing different neighbors to have varying levels of influence on a central node, enhancing the model’s performance on complex graph structures. With its attention calculation as in (1), GAT is better suited for handling heterogeneous node features and intricate dependencies [48]. In residue contact map prediction, GAT has an advantage as the interactions between different residues may vary in influence or significance.

The node features output by the graph neural network are separated into node features for each of the two input proteins. By computing the Cartesian product of these node features and concatenating the features of paired nodes, we obtain pairwise inter-residue features for all residue pairs. Fig. 3(a) illustrates the process of calculating these pairwise interresidue features. All pairwise features are then passed through two fully connected layers (FC layers) to predict the likelihood of contact between residue pairs and subsequently reshaped to generate the contact map for the two input proteins.

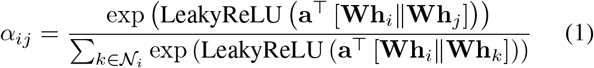

**Fig. 3.**
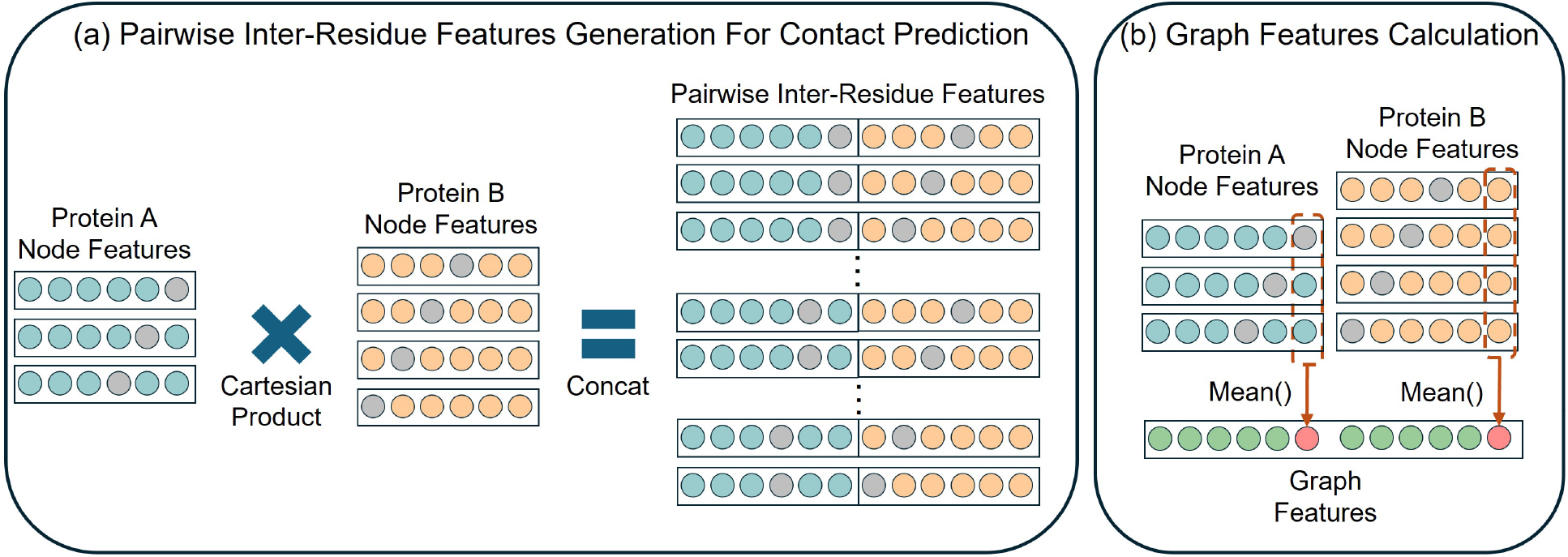
Feature processing steps for contact prediction and graph feature calculation. (a) Pairwise inter-residue features are generated by computing the Cartesian product of node features from Protein A and Protein B, followed by concatenation. (b) Graph features are calculated by applying mean pooling to node features, providing fixed-length graph representations for interaction prediction.

### D. Interaction Predictor

The Interaction Predictor uses residue pairs with a contact likelihood greater than 0.5 from the contact map predicted by the Interface Predictor to update the graph representation of the input proteins, thereby generating a complete graph representation of the potential protein complex. A GCN layer is then applied to extract information from the complete graph representation. The node features are aggregated using global mean pooling by chain to generate graph features, as shown in Fig. 3(b). Global mean pooling by chain involves applying global mean pooling separately to residues from each of the two proteins and then concatenating the resulting features to form the graph representation. The graph features are then passed through a fully connected layer (FC layer) to predict the likelihood of interaction between the two input proteins.

### E. Evaluation Methods

In this study, we compared the performance of SpatialPPI 2.0 with AlphaFold3, DeepInter, and CDPred in predicting contact maps. AlphaFold3 predictions were performed using the AlphaFold Server, selecting the top-ranked prediction results. DeepInter was obtained from its official repository and executed according to its default manual using the UniRef30 2021 03 version of the Uniclust database. CDPred was obtained from its GitHub repository, with the Uniclust database sourced from the UniRef90 01 2020 version, as specified in its manual. Since the ratio of negative to positive contacts in contact maps is highly imbalanced, with contacted residues comprising only a small fraction of the total, we used average precision (AP) as the primary evaluation metric. AP measures the overall precision-recall performance at different thresholds. In imbalanced datasets, AP focuses on the positive class predictions, mitigating the influence of the negative class. For protein-protein interaction prediction, we compared our model with state-of-the-art methods, including Topsy-Turvy, D-script, Struct2Graph, and GNN-PPI. Topsy-Turvy and D-script were obtained from their official websites and used with their pre-trained datasets for prediction. Struct2Graph and GNN-PPI were obtained from their GitHub repositories, and data were downloaded and trained following their reproducibility guidelines. The same protein structure data used in SpatialPPI 2.0 were employed for prediction. We evaluated the models using key binary classification metrics, including accuracy, precision, recall, F1 score, AP, and area under the ROC curve (AUC).

## III. Results

### A. Performance of Inter-chain Contact Maps Prediction

The performance of SpatialPPI 2.0 on the contact map prediction task was evaluated using the AP metric, due to the highly imbalanced nature of contact and non-contact residue pairs. We compared our model with AlphaFold3, DeepInter, and CDPred. Table II provides a detailed quantitative evaluation of the models in terms of AP, Precision, Recall, and F1-score. SpatialPPI 2.0 outperforms the other models across all metrics, with an AP of 0.783, precision of 0.698, recall of 0.769, and an F1-score of 0.732. AlphaFold3 ranks second, while DeepInter and CDPred achieve significantly lower scores. The higher recall and precision of SpatialPPI 2.0 illustrate its effectiveness in both identifying true contacts and minimizing false positives. The GAT layer, with its ability to handle heterogeneous node features, contributed significantly to the model’s improved performance in predicting complex protein-protein interactions. Fig. 4 illustrates the precision-recall curves for all models, showing that SpatialPPI 2.0 consistently achieves higher precision across a wide range of recall values. This demonstrates the model’s robustness in maintaining a balance between identifying true contacts and minimizing false positives. The superior performance of SpatialPPI 2.0 is attributed to its ability to capture complex interactions through the GAT layer, effectively distinguishing residue pairs that are more likely to interact. The attention mechanism used by the GAT layer allows SpatialPPI 2.0 to dynamically focus on the most important residue pairs, which contributes to the higher AP. Fig. 5 provides further insight into the model performance by visually comparing the predicted contact maps for protein pair O13978 and O36019 across different models. (a), (b), (d), and (e) represent the predictions from SpatialPPI 2.0, AlphaFold3, DeepInter, and CDPred, respectively, while (c) shows the ground truth contact map and (f) shows the distance distribution between the two proteins. This visual analysis helps to highlight the accuracy of SpatialPPI 2.0 in correctly identifying contact regions with less noise compared to other models.

**TABLE II.** Performance evaluation of models predicting the likelihood of contact between residues in two proteins, using Average Precision (AP), Precision, Recall, and F1-score.

|  | SpatialPPI 2.0 | AlphaFold3 | DeepInter | CDPred |
| --- | --- | --- | --- | --- |
| AP | 0.783 | 0.726 | 0.558 | 0.493 |
| precision | 0.698 | 0.644 | 0.51 | 0.444 |
| recall | 0.769 | 0.783 | 0.6 | 0.414 |
| F1 score | 0.732 | 0.707 | 0.551 | 0.429 |

**Fig. 4.**
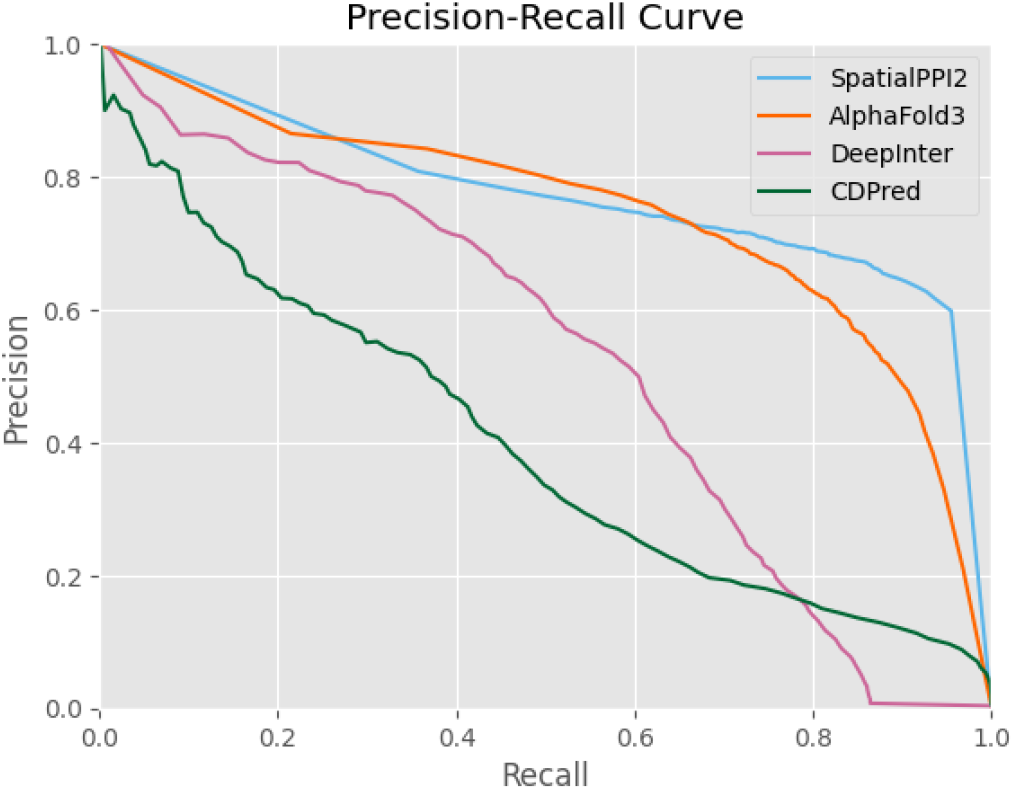
Precision-recall curves for SpatialPPI 2.0, AlphaFold3, DeepInter, and CDPred, showing the performance of each model in predicting contact maps. SpatialPPI 2.0 consistently achieves the highest precision across most recall levels, highlighting its effectiveness in distinguishing true contacts.

**Fig. 5.**
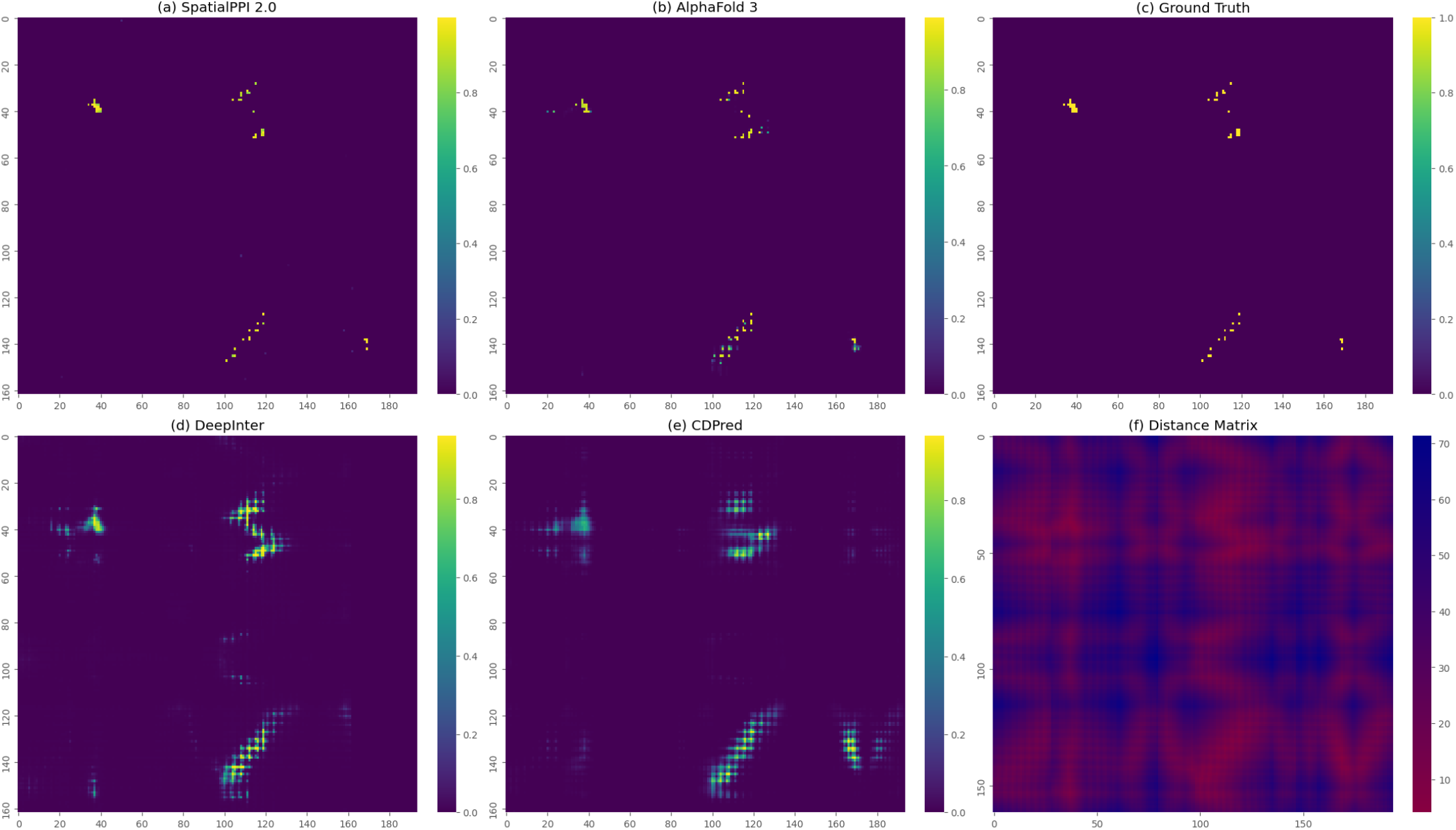
Comparison of predicted contact maps of protein pair O13978 and O36019. (a) Predicted contact map from SpatialPPI 2.0. (b) Predicted contact map from AlphaFold3. (c) Ground Truth contact map from RSCB PDB. (d) Predicted contact map from DeepInter. (e) Predicted contact map from CDPred. (f) Distribution of distance matrix from RSCB PDB. SpatialPPI 2.0 and AlphaFold3 predictions closely match the ground truth, while DeepInter and CDPred predicted the correct general regions of the contact map, they exhibit higher levels of noise.

### B. Evaluation of Protein-Protein Interaction Prediction

Table III shows the performance evaluation of five models—SpatialPPI2, Topsy-Turvy, D-script, Struct2Graph, and GNN-PPI—highlighting significant differences in their ability to predict protein-protein interactions. The receiver operating characteristic (ROC) curve in Fig. 6(a) plots the true positive rate (sensitivity) against the false positive rate, illustrating each model’s capacity to distinguish between true positive and false positive classifications. Fig. 6(b), the Precision-Recall Curve, emphasizes the models’ ability to maintain high precision while recalling true positives. SpatialPPI2 consistently outperforms the other models across multiple metrics, achieving the highest accuracy (0.826), precision (0.887), F1 score (0.811), AP (0.895), and AUC (0.896). This is further corroborated by the ROC and precision-recall curves, where SpatialPPI2 demonstrates a superior balance of true positive rate and precision across a wide range of thresholds, indicating its robust overall performance. These results indicate that SpatialPPI 2.0 can effectively maintain a balance between identifying true positive PPIs and minimizing false positives, thanks to the combination of GAT and GCN. The GAT layer, in particular, allows SpatialPPI 2.0 to assign different levels of attention to residue pairs, which enhances prediction accuracy.

**TABLE III.** Performance evaluation of models predicting the likelihood of protein-protein interactions, using Accuracy, Precision, Recall, F1 Score, AP, and AUC.

|  | SpatialPPI 2.0 | Topsy-Turvy | D-script | Struct2Graph | GNN-PPI |
| --- | --- | --- | --- | --- | --- |
| Accuracy | 0.826 | 0.788 | 0.624 | 0.793 | 0.695 |
| Precision | 0.887 | 0.831 | 0.787 | 0.745 | 0.742 |
| Recall | 0.746 | 0.723 | 0.340 | 0.890 | 0.597 |
| F1 score | 0.811 | 0.773 | 0.475 | 0.811 | 0.661 |
| AP | 0.895 | 0.842 | 0.743 | 0.846 | 0.762 |
| AUC | 0.896 | 0.866 | 0.759 | 0.876 | 0.718 |

**Fig. 6.**
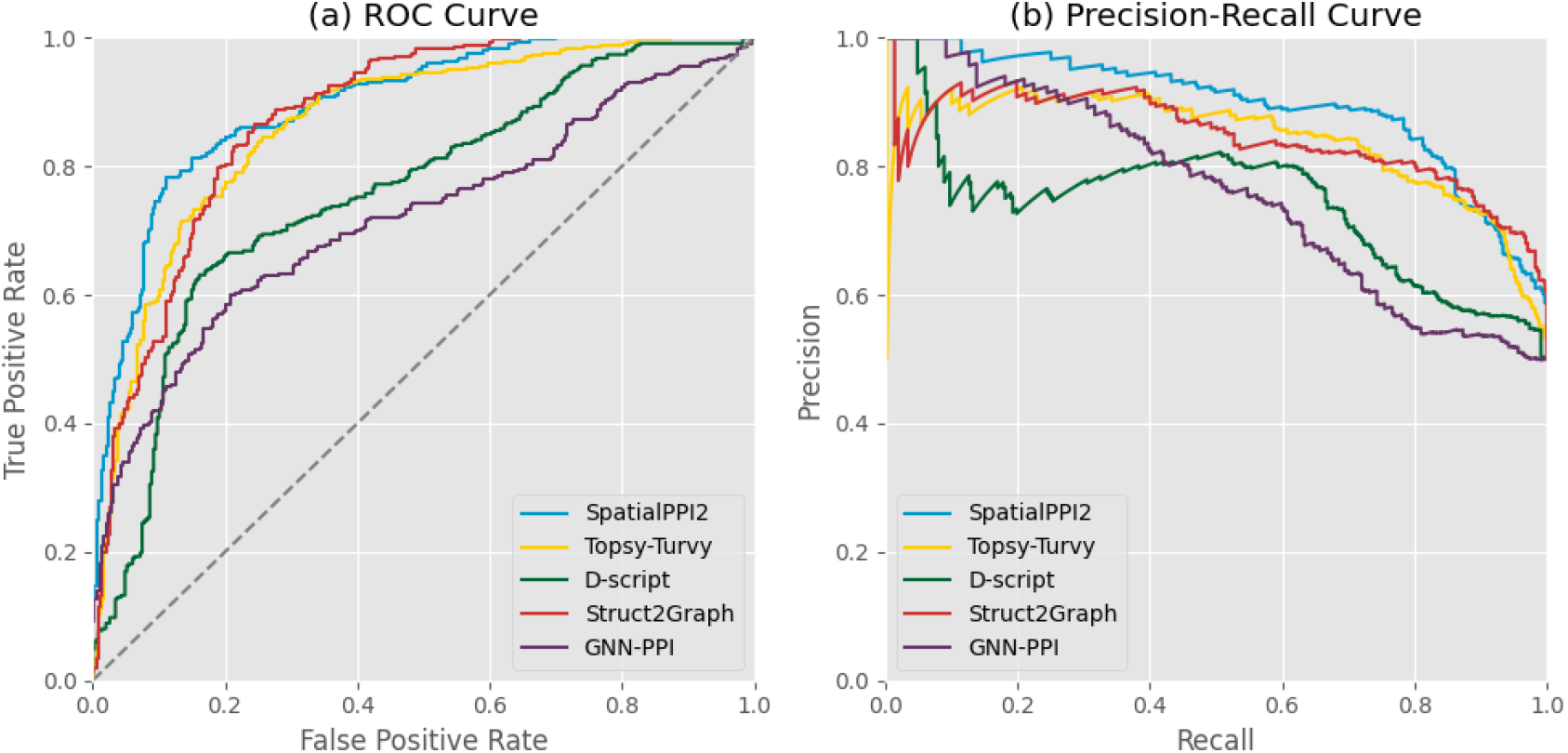
(a) Receiver Operating Characteristic (ROC) curve showing the true positive rate versus the false positive rate for SpatialPPI 2.0, Topsy-Turvy, D-script, Struct2Graph, and GNN-PPI. The ROC curve illustrates each model’s capability to distinguish between true and false positive classifications. (b) Precision-Recall curve depicting each model’s ability to maintain high precision while recalling true positive interactions. SpatialPPI 2.0 consistently demonstrates superior performance, achieving a high balance between recall and precision across multiple thresholds.

Struct2Graph also performs well, particularly in terms of recall (0.890) and F1 score (0.811), and maintains competitive performance in both the ROC and precision-recall curves, suggesting its utility in scenarios where high recall is essential. Topsy-Turvy demonstrates a solid balance between precision (0.831) and AUC (0.866), making it a competitive option for tasks requiring high precision. However, the D-script performs poorly, with notably low recall (0.340) and F1 score (0.475), as well as degraded performance in the precision-recall curve, indicating significant limitations in identifying positive samples. GNN-PPI, while exhibiting moderate precision (0.742), suffers from lower recall (0.597) and AUC(0.718), as reflected in its inferior performance in both curves, suggesting that further refinement is needed to improve its overall discriminatory power.

### C. Predicting with AlphaFold

AlphaFold does not always successfully determine whether the input proteins can interact. In some cases, even when the input proteins cannot physically interact, AlphaFold produces a tightly bound protein complex. Fig. 7 presents AlphaFold3’s prediction for the protein pair P0C8E7 and P13501, taken from Negatome 2.0, which are experimentally verified to lack interactions. In (a), the contacted residues between the two proteins are highlighted, revealing substantial interaction regions despite the absence of interaction. In (b), the predicted local Distance Difference Test (pLDDT) score is shown, representing the prediction confidence; notably, the interacting regions exhibit the highest confidence level (¿90). Panel (c) displays the Predicted Aligned Error (PAE) for the predicted structure by AlphaFold3. PAE estimates the error in the relative position and orientation between two residues in the predicted structure, with higher values indicating greater uncertainty and, thus, lower confidence. It can be observed that the model also shows a low PAE for the interacting region.

**Fig. 7.**
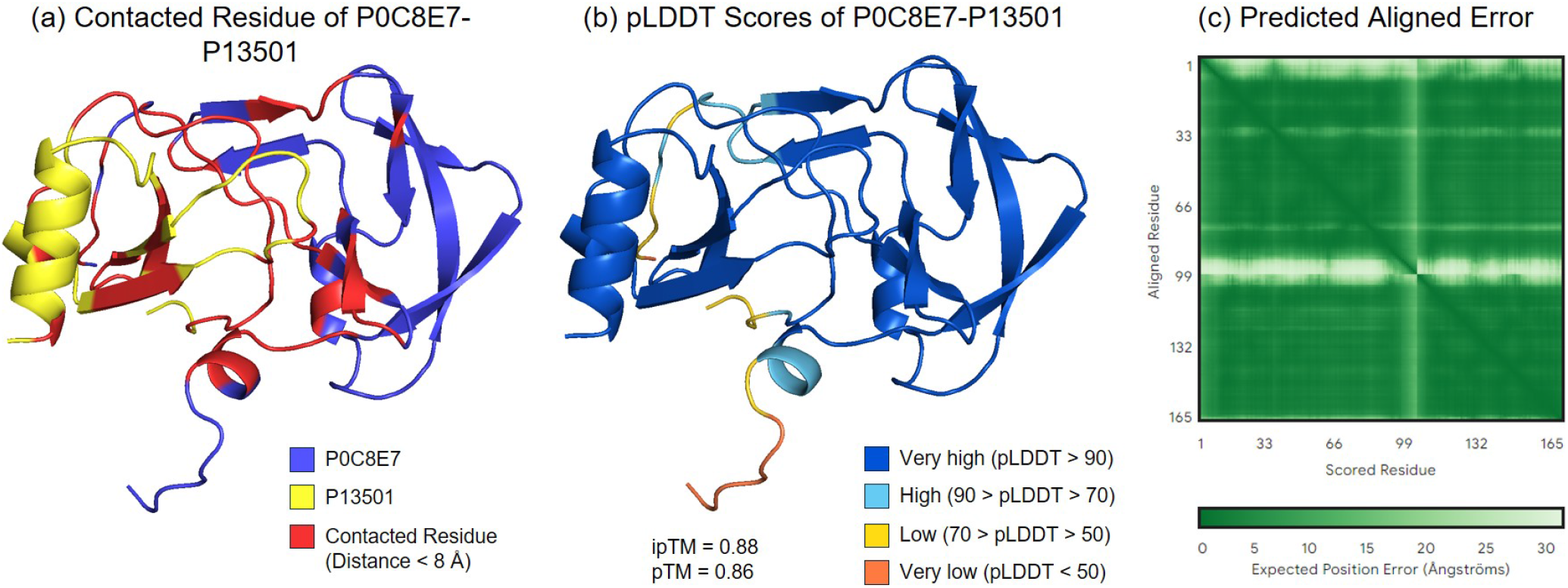
AlphaFold3 predictions for the protein pair P0C8E7 and P13501, which are experimentally verified to lack interactions. (a) Contacted residues between the two proteins predicted by AlphaFold3, showing substantial predicted interaction regions. (b) pLDDT score represents prediction confidence, with higher scores indicating high confidence in the structure. (c) Predicted Aligned Error (PAE) estimating error in the relative position and orientation between residues, with lower PAE values indicating higher confidence in the predicted structure.

Nevertheless, AlphaFold provides a potential solution to address the scarcity of protein structure data. In this study, structural data from the AlphaFold Database was used as input to SpatialPPI 2.0, and the results were compared against predictions based on authentic protein structures from PINDER gathered from the RCSB PDB database. Additionally, predictions were made using complexes directly predicted by AlphaFold3 as inputs for SpatialPPI 2.0’s Interaction Predictor. The results are summarized in Fig. 8 and Table IV. Predictions using structures from the AlphaFold Database showed a slight decrease in accuracy compared to those using real structural data but still demonstrated satisfactory overall performance. Interestingly, predictions using AlphaFold3 Predicted Complexes were comparable to those using authentic data. The ROC curve in Fig. 8(a) shows that the predictive capacity for distinguishing true and false positives using AlphaFold3 Predicted Complexes is similar to that using authentic data. The Precision-Recall curve in Fig. 8(b) indicates minor instability when using predicted structures as input. Overall, these tests demonstrate that SpatialPPI 2.0 remains robust when utilizing predicted structure data, offering a promising solution for predicting interactions between proteins for which experimentally confirmed structural information is unavailable.

**TABLE IV.** Performance Evaluation of Protein-Protein Interaction Prediction Using Structures from PINDER, AlphaFold Database, and AlphaFold3 Predicted Complexes.

|  | PINDER | AlphaFold DB | AlphaFold3 Complexes |
| --- | --- | --- | --- |
| Accuracy | 0.826 | 0.762 | 0.840 |
| Precision | 0.887 | 0.867 | 0.873 |
| Recall | 0.746 | 0.620 | 0.795 |
| F1 Score | 0.811 | 0.723 | 0.833 |
| AP | 0.895 | 0.849 | 0.880 |
| AUC | 0.896 | 0.858 | 0.892 |

**Fig. 8.**
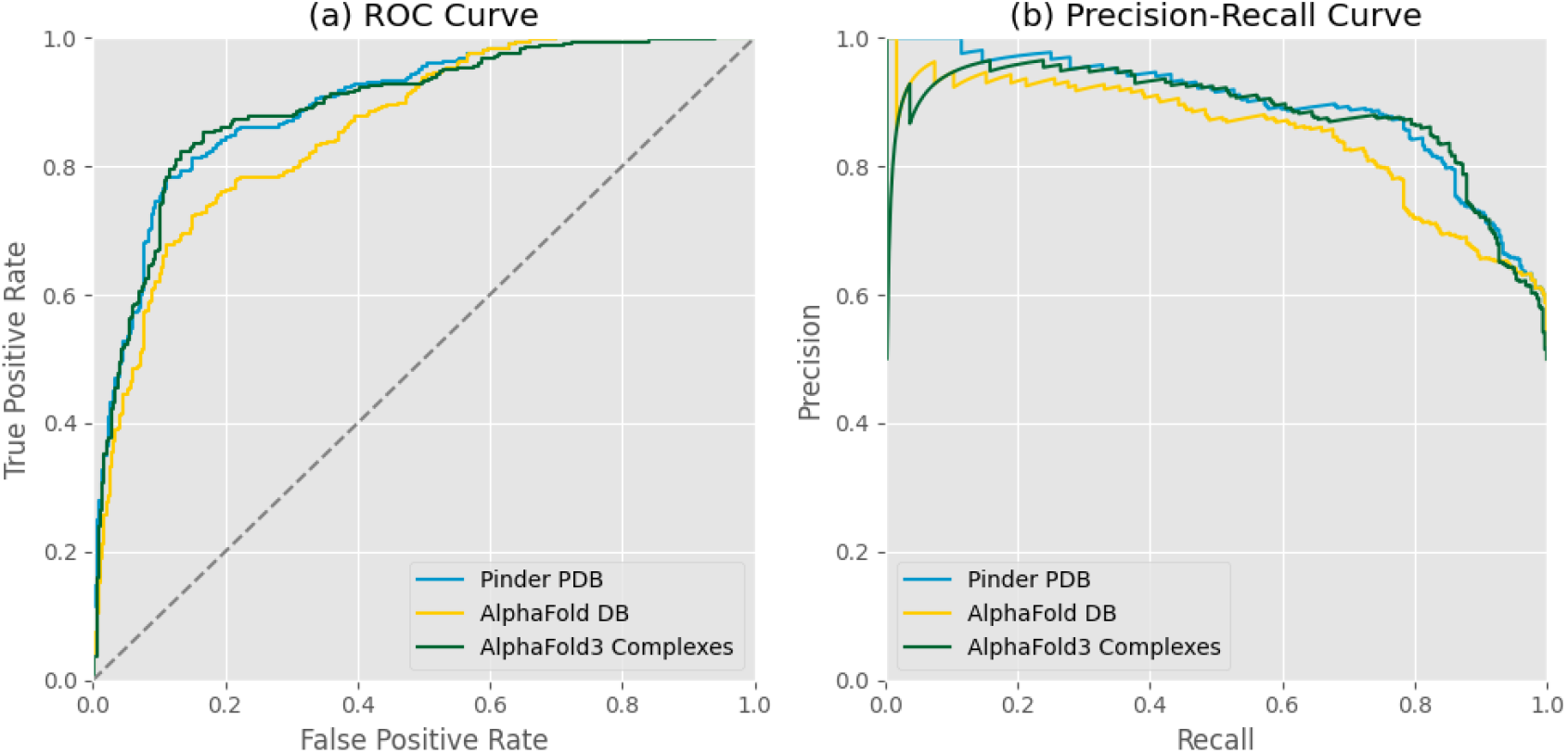
Receiver Operating Characteristic (ROC) curve showing the true positive rate versus false positive rate for predictions using AlphaFold3 predicted complexes, AlphaFold database, and PDB from PINDER as inputs. The ROC curve illustrates that the predictive capacity for distinguishing true and false positives using AlphaFold3 predicted complexes closely matches that using authentic data. (b) The precision-recall curve shows slight instability when using predicted structures as input, indicating minor discrepancies in precision at different recall levels.

## Conclusion and Discussion

SpatialPPI 2.0 presents significant advancements in proteinprotein interaction (PPI) prediction over its predecessor. Unlike the original SpatialPPI, SpatialPPI 2.0 predicts interaction interfaces as intermediate steps to enhance the prediction of interactions between isolated proteins. This two-step process—first predicting contact maps, then assessing overall interaction—improves specificity by focusing on relevant regions of protein structures, reducing false positives.

The use of GAT allows dynamic assignment of importance to residues, capturing varying roles in protein interactions, while GCN contributes to capturing both local and global structural features. The synergy between GAT and GCN provides a more comprehensive representation of protein complexes.

SpatialPPI 2.0 also incorporates ProtBert, a language modelbased feature extractor, which provides rich contextual biochemical information. This allows the model to predict interactions involving novel or engineered proteins, which is highly beneficial in synthetic biology and drug discovery.

Furthermore, SpatialPPI 2.0 demonstrates robustness when using AlphaFold-predicted structures, achieving comparable performance to predictions using experimental data. This ability to leverage structural predictions broadens its applicability in cases where experimental structures are unavailable.

In summary, SpatialPPI 2.0 achieves superior performance by integrating interaction interface prediction, advanced graph neural networks, and language model-derived features. Its unique multi-step approach and robustness with predicted structures make it a powerful tool for advancing the study of protein interactions, aiding in drug discovery, synthetic biology, and the understanding of complex biological processes.

## Acknowledgment

This work was financially supported by JST FOREST (JPMJFR216J), JSPS KAKENHI (JP23H03496), and AMED BINDS (JP24ama121026). The computational experiments were performed using the TSUBAME 4.0 supercomputer at the Institute of Science Tokyo.

## Notes

### Competing Interest Statement

The authors have declared no competing interest.

